# Intraperitoneal transfer of wild-type bone marrow cells in the *Csf1r* knockout rat repopulates resident tissue macrophages without contributing to monocytopoiesis

**DOI:** 10.1101/2023.02.19.529164

**Authors:** Anuj Sehgal, Dylan Carter-Cusack, Sahar Keshvari, Omkar Patkar, Stephen Huang, Kim M. Summers, David A. Hume, Katharine M. Irvine

**Author notes:** These authors contributed equally (Anuj Sehgal, Dylan Carter-Cusack). Joint senior authors (David A. Hume, Katharine M. Irvine).

## Abstract

Homozygous null mutation of the macrophage colony-stimulating factor receptor (*Csf1r*) gene in rats leads to the loss of most tissue macrophage populations and has pleiotropic impacts on postnatal growth and organ maturation leading to mortality by 8-12 weeks of age. The phenotype of the *Csf1r* knockout (*Csf1rko*) can be reversed by intraperitoneal transfer of wild-type bone marrow cells (BMT) at weaning. Here we used a *Csf1r*-mApple transgenic reporter, which is expressed in neutrophils and B cells as well as monocytes and macrophages, to track the fate of donor-derived cells. Following BMT into *Csf1r* recipients, wild-type mApple^+ve^ cells restored IBA1^+^ tissue macrophage populations in every tissue donor-derived cells also completely replaced recipient macrophages in organs such as spleen, lung and liver that were only partly macrophage-deficient in the *Csf1rko*. However, monocytes, neutrophils and B cells in bone marrow, blood and lymphoid tissues remained of recipient (mApple^-ve^) origin. An mApple^+ve^ cell population expanded in the peritoneal cavity and invaded locally in the mesentery, fat pads, omentum and diaphragm. One week after BMT, distal organs contained foci of mApple^+ve^, IBA1^-ve^ immature progenitors that appeared to proliferate, migrate and differentiate locally. We conclude that rat bone marrow contains progenitor cells that are able to restore and maintain all tissue macrophage populations in a *Csf1rko* rat directly without contributing to the bone marrow progenitor or blood monocyte populations.

## Introduction

Resident macrophages are an abundant cell population in every major organ. They share a regular distribution associated with endothelial and epithelial basement membranes, stellate morphology, a core transcriptome and basic endocytic functions but they also adapt specifically to each tissue environment [1-4]. The differentiation, survival and maintenance of macrophages in all vertebrates is controlled primarily by two ligands, colony stimulating factor 1 (CSF1) and interleukin 34 (IL34), each of which signals through the CSF1 receptor (CSF1R) (reviewed in [5-9]. Macrophages are abundant throughout the embryo, produced first in the yolk sac and then in fetal liver, and actively phagocytose apoptotic cells [10, 11]. Surprisingly, the almost complete absence of macrophages throughout gestation in *Csf1r* hypomorphic mutant mice [12, 13] or *Csf1r* knockout (*Csf1rko*) rats [14] has little apparent effect on development of the embryo. The main phenotypic impacts of macrophage deficiency occur in the postnatal period, with ventricular enlargement in the brain (hydrocephalus), growth retardation, failure of skeletal development and osteopetrosis and delayed maturation of major organs, notably the liver [5, 14-16]. Although the *Csf1rko* rats are globally macrophage-deficient, CSF1R expression is not required for tissue-specific adaptation. Expression profiling of liver [14] indicated the residual Kupffer cells (KC) still expressed KC-specific transcripts such as *Clec4f, Cd5l* and *Vsig4*.

In inbred *Csf1rko* mice, hydrocephalus leads to pre-weaning lethality [5, 15] whereas inbred *Csf1rko* rats can survive 7-10 weeks despite significant ventricular enlargement [14, 16, 17]. Similar brain and skeletal pathologies observed in rats also present in rare individuals with homozygous recessive *CSF1R* mutations in humans [18-21]. Bennett *et al*.[22] reported that populations of brain microglia-like cells and the pre-weaning lethality of the *Csf1rko* in C57BL/6J mice could be rescued with around 50% success by intra-peritoneal transfer of wild-type (WT) bone marrow cells at birth. Beers *et al*. [23] reported similar outcomes following neonatal transfer of WT marrow in PU.1 (*Spi1*) knockout mice, which also lack embryonic macrophage populations. These two studies inferred without direct evidence that WT donor cells migrate to the bone marrow and reconstitute normal monocytopoiesis in the recipients leading to repopulation of tissue-resident macrophages derived from blood monocytes. However, monocyte production *per se* is not CSF1R-dependent [24]. *Csf1rko* rats have a selective loss of the non-classical monocyte subset (His48^low^/CD43^high^, equivalent to Ly6C^low^ in mice, CD16^high^ in human) which constitutes the majority of monocytes in this species [14, 25]. *Csf1rko* rats have a significant granulocytosis and B cell-deficiency but despite the apparent loss of hematopoietic island macrophages in fetal liver and bone marrow, they are not anemic or leukopenic.

The *Csf1rko* rat was generated and characterised on the dark agouti (DA) inbred background [16]. On this background, the pleiotropic *Csf1rko* phenotypes were almost completely reversed by intraperitoneal transfer of BM cells from congenic WT animals as late as weaning [14]. The tissue macrophage populations were restored but the rats remained monocyte-deficient and the bone marrow compartment remained CSF1-resistant, despite the reversal of the osteopetrosis and full restoration of cellularity in the marrow cavity [14]. Hence, the mechanism of tissue macrophage repopulation was not evident. Since the *Csf1rko* rats are not entirely macrophage deficient, it remains possible that restoration of osteoclasts and bone marrow cellularity enables repopulation of some or all populations by recipient cells. It is also unclear whether donor cells contribute to the restoration of normal granulocyte and B cell populations.

To investigate the mechanism, we have taken advantage of a *Csf1r*-mApple reporter transgene [25]. This reporter transgene was generated on an outbred Sprague-Dawley (SD) background whereas the *Csf1rko* is on the inbred dark agouti (DA) background. We previously intercrossed these lines and demonstrated on the mixed background that the *Csf1rko* leads to profound loss of *Csf1r*-mApple^+^ macrophages in most tissues [14]. The transgene is also expressed in granulocytes (which express *Csf1r* mRNA, but not protein; [26]) and in some B cells, so visualization of the macrophage deficiency in hematopoietic/lymphoid tissues using this reporter is compromised. However, this expression enables analysis of WT marrow contribution to recovery of non-macrophage (neutrophil and B cell) populations in bone marrow transfer (BMT) recipients. Here we have generated an inbred SD line segregating the *Csf1rko* allele and *Csf1r-*mApple reporter. Our analysis using this model indicates that donor cells proliferate in the peritoneal cavity, and immature progenitors disperse through lymph and blood to seed and repopulate every tissue macrophage population. Donor cells did not contribute to any circulating blood cell lineage. We suggest that bone marrow contains a committed macrophage progenitor population with high proliferative potential that can entirely repopulate the mononuclear phagocyte system without a monocyte intermediate.

## Results

### The *Csf1rko* phenotype is maintained in an inbred Sprague-Dawley-derived line

The phenotypic impact of the *Csf1rko* in mice is strongly influenced by genetic background and is pre-weaning lethal in inbred lines such as C57BL/6J [5]. To avoid this issue, the initial analysis of the rat *Csf1rko* was carried out on a mixed genetic dark agouti (DA)/outbred Sprague-Dawley (SD) genetic background [16]. The *Csf1rko* allele was subsequently crossed for 5 generations to a commercial outbred SD line and separately to a pure inbred DA background. On the DA background, wild-type bone marrow transfer (BMT) at weaning restored tissue macrophage populations and rescued the phenotype [14]. The transgenic *Csf1r-*mApple (*Csf1r*^mApple^) reporter transgenic line was generated and analysed originally on the outbred SD background [25]. The same reporter in mice was expressed in monocyte-macrophages, neutrophils and B cells [27]. To enable the transfer of *Csf1r*^mApple^ bone marrow into *Csf1rko* recipients, the 2 SD lines were crossed, and then inbred to retain both the mutant allele and reporter on the same background. Despite the inbreeding, the SD line retained large litter sizes which expedited generation of *Csf1rko* pups from heterozygous matings. The inbred *Csf1rko*^SD^ pups grew more rapidly than the inbred DA *Csf1rko* but nevertheless were severely osteopetrotic and showed the same growth arrest by around 7-8 weeks of age (**Figure 1A**). As in the DA background, the reduced somatic growth was associated with very low serum IGF1 levels at 3 weeks of age (**Figure 1B**). Serum IGF1 has not been measured on the SD background at later time points, but IGF1 remained low in older *Csf1rko* rats on the DA background [14].

**Figure 1.**
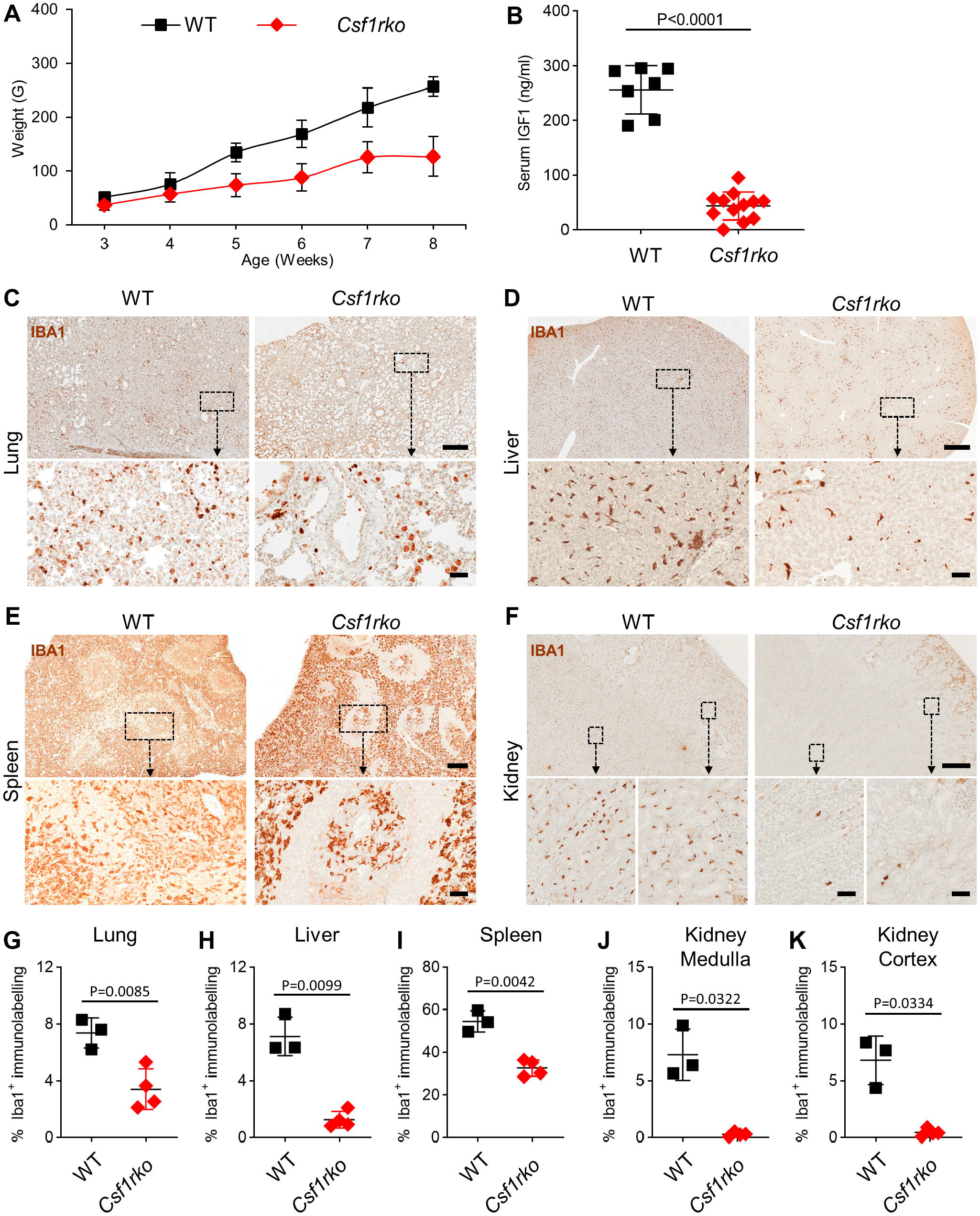
Postnatal growth retardation and tissue macrophage deficiency in *Csf1rko* rats. (A) Time course of weight gain of WT^SD^ (n=12) and *Csf1rko*^SD^ (n=19) rats. (B) IGF1 serum concentrations in WT^SD^ (n=7) or *Csf1rko*^SD^ (n=12) rats at 3 weeks of age. (C–F) IBA1 immunolabelling in lung (C), liver (D), spleen (E) and kidney (F) at 3 weeks of age. Scale bars in main images for (C), (D) and (F) are 500μm. Scale bar for (E) is 250μm. All magnified scale bars at 50μm. (G–K) Quantification of percentage IBA1 immunolabelling in field of view of lungs (G), liver (H), spleen (I), kidney medulla (J) and kidney cortex (K). Each data point represents mean of 4 fields of view analyzed per animal (n=3-4 per genotype). Two-tailed unpaired Student’s t-test.

The loss of macrophages in tissues of the *Csf1rko* rat can be visualised using IBA1 as a marker [14]. On the SD genetic background, there were some residual IBA1^+^ cells detected in lung and liver in juvenile (3 weeks) *Csf1rko* rats whereas the kidney was entirely depleted and the overall population in the spleen was reduced by around 50% (**Figure 1C-F**; quantified in **Figure 1G-K**). The initial description of the *Csf1rko* rat on a mixed background noted the selective loss of marginal zone macrophages and associated transcripts (e.g. *Siglec1, Cd209f*) in the spleen whereas the red pulp population was less affected [16]. The selective loss of IBA1^+^ macrophages from the marginal zone in the *Csf1rko*^SD^ is evident in **Figure 1E**. Interestingly, the population of IBA1^+^ cells in the periarteriolar lymphoid sheath (PALS) was not depleted and indeed appeared more prominent. There was also a consistent increase in apparent IBA1 staining intensity in individual macrophages of the red pulp in the *Csf1rko* although the proportional area of staining was reduced.

### The *Csf1r*-mApple reporter enables detection of donor-derived cells in *Csf1rko* recipients

We transferred wild-type (WT) *Csf1r*-mApple bone marrow cells (BMT) to congenic *Csf1rko*^SD^ recipients by IP injection at 2 weeks of age (**Figure 2A**). By 2-3 weeks post BMT the growth rate of the recipients accelerated and diverged from *Csf1rko* non-transplanted controls (**Figure 2A**). The low serum IGF1 levels seen at 3 weeks **(Figure 1B)** were restored to wild-type levels in the long-term transplant recipients (**Figure 2B**). The initial cohort of BMT rats generated sexually mature male and female animals that were able to produce litters when mated to heterozygous animals to generate litters in which 50% of progeny were *Csf1rko*.

**Figure 2.**
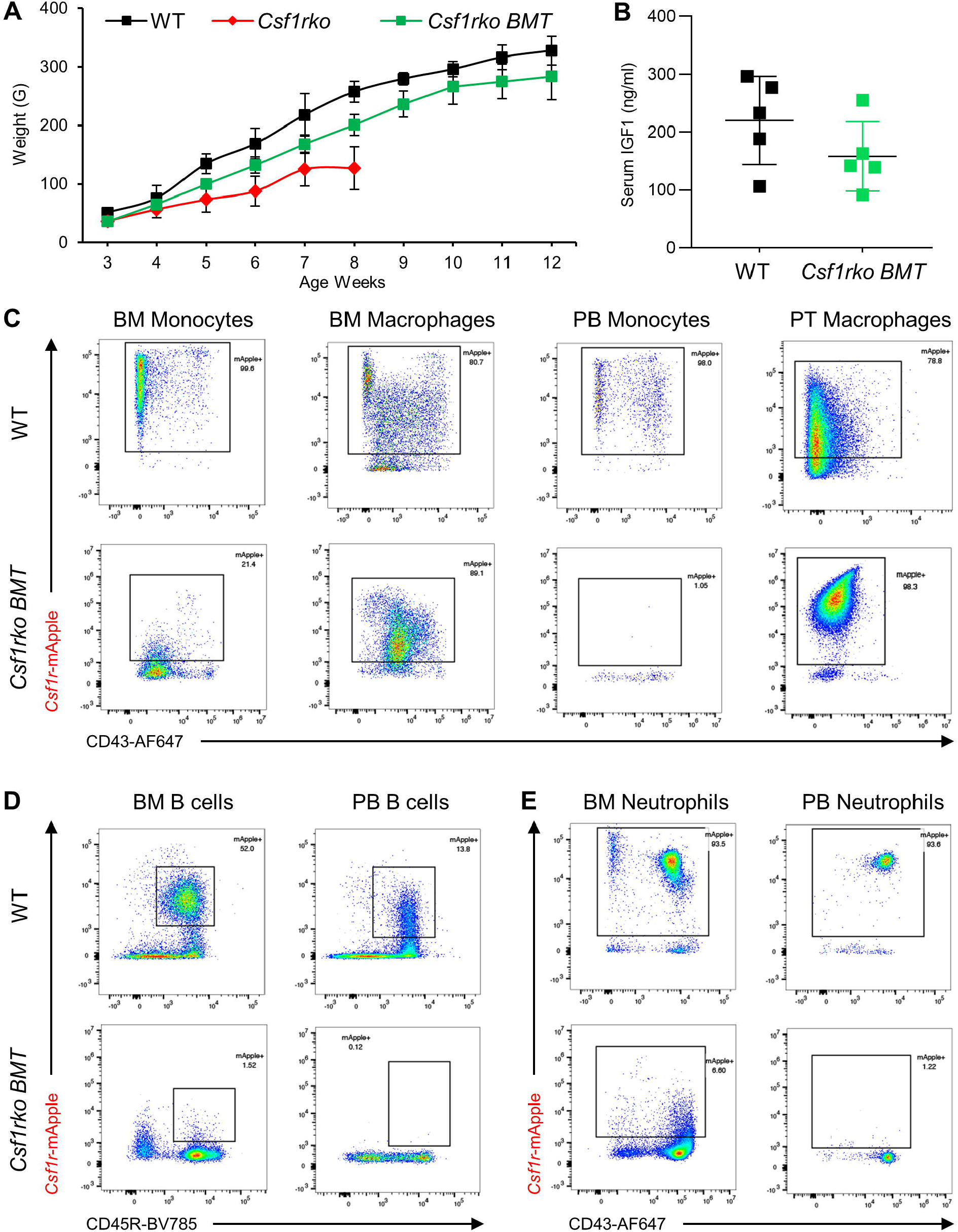
Intraperitoneal transfer of WT *Csf1r*^mApple^ bone marrow cells restores growth without contributing to recipient hematopoiesis. *Csf1rko* rats (2 females and 3 males) received 2×10^7^ WT *Csf1r*^mApple^ female bone marrow (BM) cells by intraperitoneal injection (BMT) at 2 weeks of age. (A) Time course of weight gain in BMT recipients. The growth curves of WT and *Csf1rko* from Figure 1A are shown for comparison. (B) IGF1 serum concentrations in WT littermates and *Csf1rko*-BMT rats and were analyzed a minimum of 16-weeks post-transplant. (C) Flow cytometry analysis of BM monocytes, putative BM macrophages, peripheral blood (PB) monocytes and peritoneal (PT) macrophages in *Csf1r*^mApple^ donor and *Csf1r*-BMT recipient rats (>16 weeks of age). (D–E) Flow cytometry analysis of BM and peripheral blood B cells (D) and neutrophils (E) in *Csf1r*^mApple^ donor and *Csf1r*-BMT recipient rats. Gating strategies are shown in Supplementary Figure 1. Profiles in Panels C-E are representative of the 5 BMT recipients and donors of similar age.

**Figure 2C-E** shows Flow Cytometry analysis of peripheral blood, peritoneal cells and bone marrow cells in long-term (>12 weeks) BMT recipients compared to the wild-type *Csf1r*^mApple^ donors. The gating strategies for this analysis are provided in **Figure S1**. Analysis of bone marrow is complicated by the phenomenon of macrophage fragmentation, which leads to unrelated cells being coated with macrophage-derived remnants [28]. In the WT donor marrow and blood, mApple was detected in monocytes (CD172^lo^/His48^+^/SSC^lo^), macrophages (CD172^+^/CD4^+^/CD43^+^), neutrophils (His48^hi^/SSC^hi^) and B cells (CD45R^+^). As noted previously [25], in the bone marrow the majority of monocytes are classical (His48^hi^, CD43^low^) whereas this distribution is reversed in the blood (**Figure S1A/B**). In the long term BMT recipients, there was very low detection of mApple in monocytes, neutrophils and B cells in marrow. This was most likely due to macrophage remnant contamination [28] since despite the lack of expression in monocytes and progenitors, the BM resident macrophages were clearly donor-derived and mApple^+ve^. Indeed, the marrow macrophage population in the BMT rats appeared more uniform and strongly mApple^+ve^/CD43^+ve^ compared to the donor (**Figure 2C**).

The lack of contribution of donor-derived cells to marrow hematopoiesis was supported by analysis of the blood. The rescued *Csf1rko* rats remained monocyte-deficient as described previously in the DA line [14] and the residual His48^high^CD43^low^ classical monocytic cells were of recipient origin (mApple^-ve^) **(Figure 2C)**. The circulating neutrophil or B cell populations recovered to normal levels observed in age-matched controls but without any detectable contribution from mApple^+ve^ donor cells (**Figure 2D/E)**. In the peritoneal cavity, there was complete restoration of a typical large resident macrophage population (CD172^+^/CD4^high^/CD43^-ve^) entirely of donor (mApple^+ve^) origin (**Figure 2C**).

A series of wholemount images from long-term (>12 weeks post-BMT) *Csf1rko* BMT recipients showed engraftment and complete repopulation of tissue resident macrophage populations in every organ including microglia-like cells in the brain cortex by donor-derived mApple^+ve^ cells **(Figure 3**). Since donor cells do not contribute to granulocyte and B cell pools, the reporter transgene enables visualisation of resident CSF1R^+^ macrophages in lymphoid tissues (spleen and lymph-node). **Figure 3** highlights the similar relative abundance and the striking regular distribution of mApple^+ve^ cells in every tissue, including several (thyroid, pancreas, endometrium, epididymis, tendon, eye sclera, mammary gland, brown adipose, tongue) that have not been analysed previously. In overview, the distribution and morphology of resident macrophages in the rescued *Csf1rko* rat is similar to the donor and to previous analysis of a FusionRed knock-in reporter in the mouse *Csf1r* locus [26].

**Figure 3.**
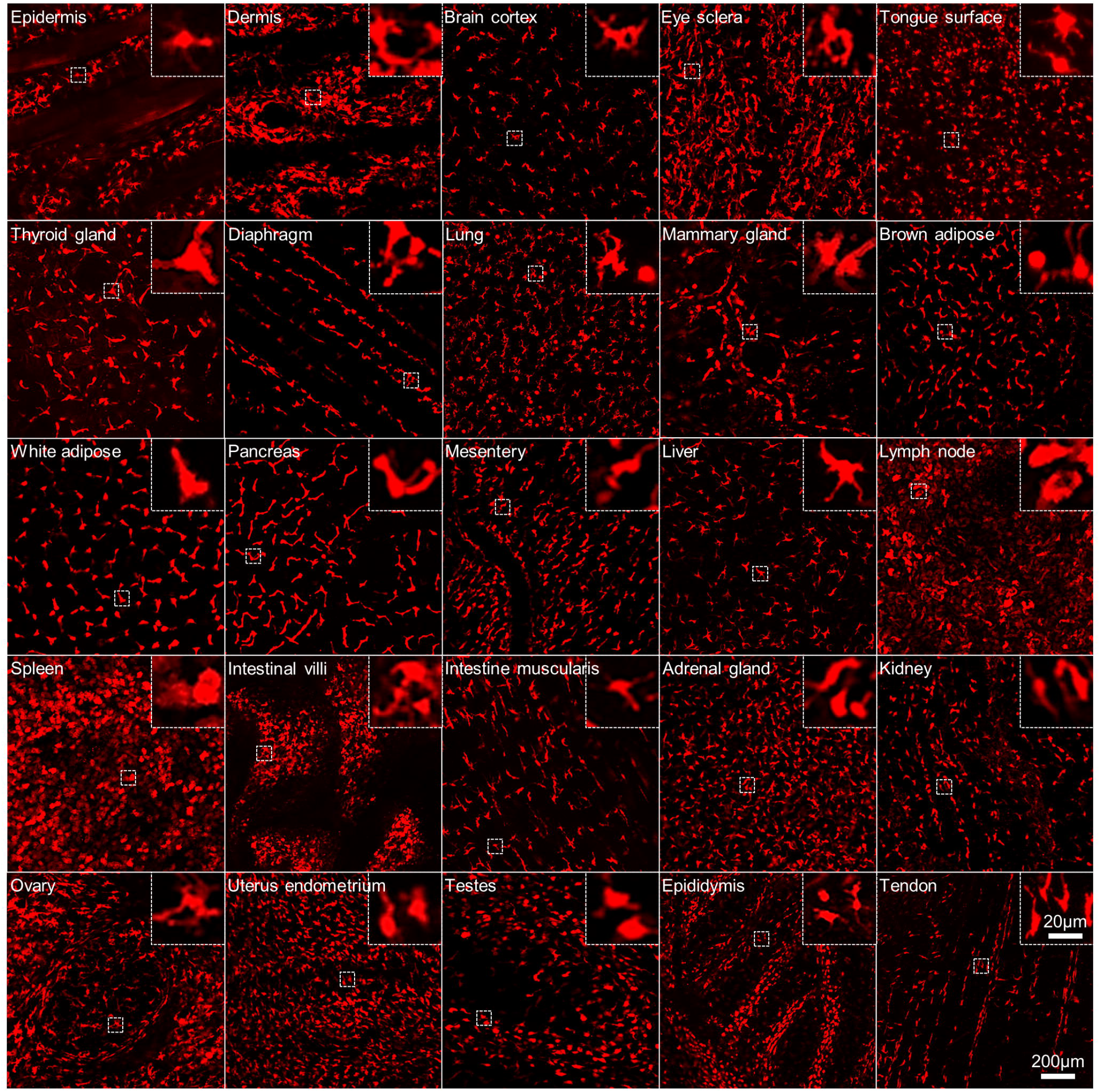
Spatial localization of *Csf1r*^mApple^ donor cells in rescued *Csf1rko* BMT recipients. *Csf1rko* rats (2 females and 3 males) received 2×10^7^ WT *Csf1r*^mApple^ female bone marrow (BM) cells by intraperitoneal injection (BMT) at 2 weeks of age and were analyzed a minimum of 16-weeks post-transplant. Tissues including skin (epidermis and dermis), brain (cortex imaged), eyes (sclera), tongue, thyroid gland, diaphragm, lung, mammary glands, brown and white adipose, pancreas, mesentery membrane, liver, mesenteric lymph nodes, spleen, small intestine (villus tip and muscularis imaged), adrenal glands, kidney (cortex), ovary, uterus (endometrium imaged), testes, epididymis, and tendon (peroneal), were extracted for wholemount immunofluorescence imaging. Images are representative of the five animals analyzed (except for reproductive tissues which represent 2 females and 3 males). No fluorescent signal is detected in any tissue in non-transgenic WT rats or in untransplanted *Csf1rko* rats. Main image scale bars = 200μm, inset 20μm. All tissues are scaled to the same settings.

**Figure 1** shows that *Csf1rko* rats were not entirely IBA1^+^ macrophage-deficient in lung, liver and spleen. In liver, around 30% of Kupffer cells remained, and in lung and spleen the population of IBA1^+^ cells was reduced by 50% at 3 weeks of age. Based upon the extensive repopulation in **Figure 3**, donor cells expressing CSF1R appear to have a competitive advantage and supplant recipient cells in locations where they are not entirely deficient in the *Csf1rko*. To confirm this interpretation, selected tissues of long term-*Csf1rko* BMT recipients were stained to detect both IBA1 and mApple (using anti-red fluorescent protein (RFP) antibody). **Figure 4** shows representative images of lung, spleen, liver, kidney and adrenal glands and quantitation of double-positive donor cells. In most locations all of the IBA1^+^ cells were mApple^+ve^ indicating complete repopulation and/or replacement by donor-derived cells. The one exception was observed in the spleen. In this organ, the marginal zone was repopulated by donor cells and the red pulp macrophages were also entirely of donor origin. However, the IBA1^+ve^ cells of the PALS, which were not depleted in the *Csf1rko* (**Figure 1**) and which express mApple in the *Csf1r*^mApple^ donor [25] remained of recipient origin.

**Figure 4.**
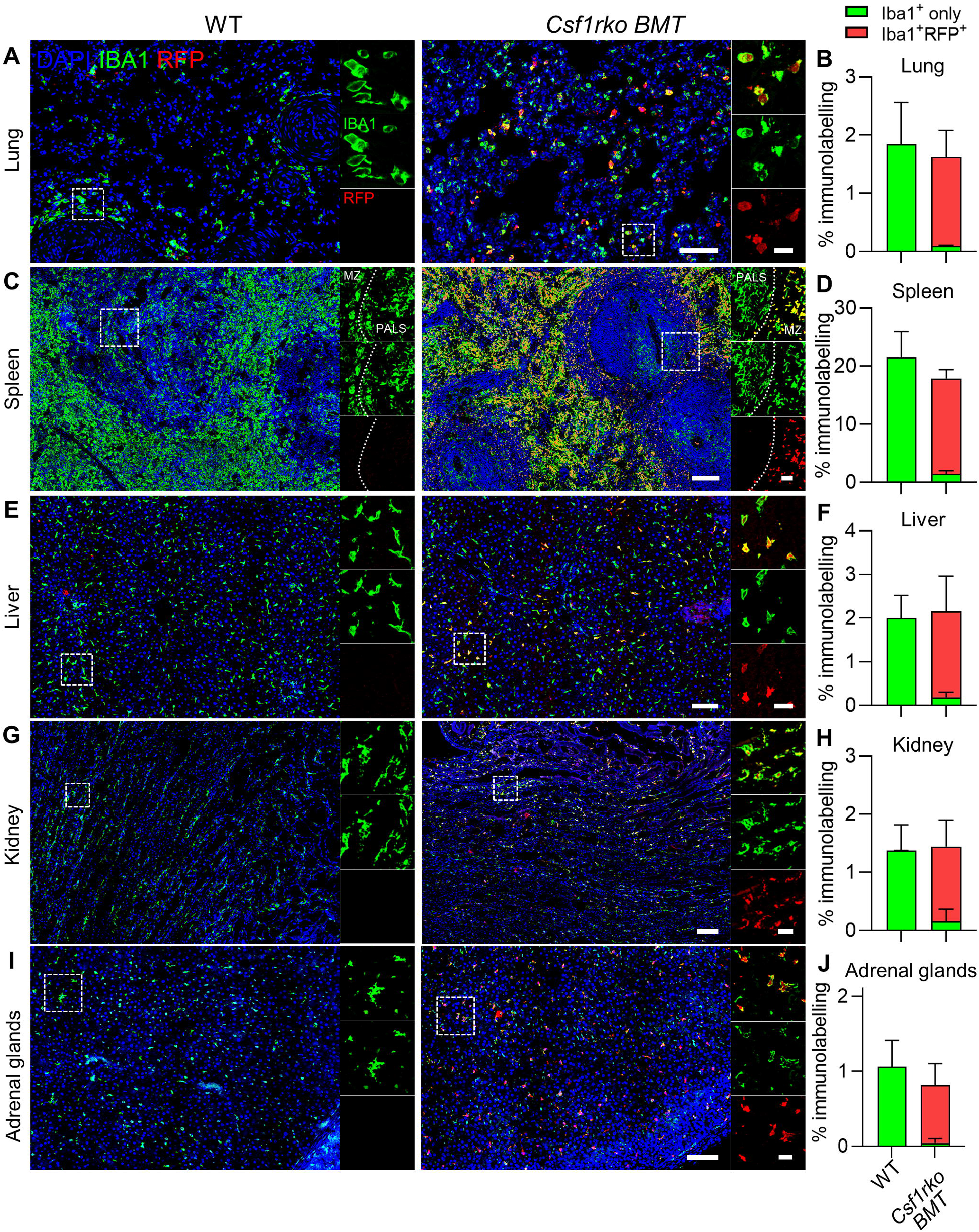
Donor-derived *Csf1r*^mApple^ cells restore and replace tissue resident macrophages in transplanted *Csf1rko* rats. *Csf1rko* rats (2 females and 3 males) received 2×10^7^ WT *Csf1r*^mApple^ female bone marrow (BM) cells by intraperitoneal injection (BMT) at 2 weeks of age and were analyzed a minimum of 16-weeks post-transplant. 5 μm PFA-fixed frozen tissue sections of lung (A), spleen (C), liver (E), kidney (G) and adrenal glands (I) from WT littermates and *Csf1r-BMT* rats were stained for IBA1 (green), RFP (red; for mApple) and DAPI (blue). In the inset panels on right hand side of main images, top panel shows merge of IBA1 and RFP, middle panel shows IBA1 and bottom panel shows RFP. In spleen images, PALS = periarteriolar lymphoid sheath; MZ = marginal zone. All tissues are scaled to the same settings so that main image scale bars = 100 μm and inset image scale = 20 μm. Morphometric analysis of tissues in corresponding graphs (B,D,F,H,J) shows total IBA1 immunolabelling to indicate total tissue resident macrophage pool and proportion of RFP^+^ immunolabelling (overlapping or within 4 pixels) indicated in the red portion of the bars. Data are derived from at least 2 sectional depths and 4 different fields of view per section from 5 animals per group. All total IBA1^+^ immunolabelling is non-significant whilst all RFP^+^ co-immunolabelling is *P*<0.0001 by two-tailed unpaired Student’s t-test (not indicated on graphs).

### Donor-derived bone marrow cells traffic from the peritoneum and repopulate tissue macrophages exclusively in *Csf1rko* recipients

The previous section showed that WT cells injected into the peritoneal cavity repopulated all the tissues of the body without contributing to bone marrow hematopoiesis or sustained recovery of circulating monocytes. To dissect the mechanism and to determine whether engraftment required a vacant peritoneal niche, *Csf1r*^mApple^ bone marrow cells were transferred into *Csf1rko, Csf1r*^*+/+*^ (WT) and *Csf1r*^+/-^ (Het) recipients and the peritoneal cells were harvested 1, 2 or 4 weeks later (**Figure S2**). In *Csf1r*^+/-^ rats, the receptor is expressed at 50% of the wild-type level [16]. We speculated that in heterozygous mutant animals, WT cells might compete for locally-available CSF1, at least in the peritoneal cavity. In the donor marrow, granulocytes (CD172a^Low^/HIS48^+^) and B cells (CD45R^+^) comprise the majority of mApple^+ve^ cells and the monocytes are predominantly His48^Hi^/CD43^low^ (**Figure S3A**). After 1 week in the *Csf1rko* recipients, the vacant peritoneal niche was repopulated and >90% of CD45^+ve^ lavage cells were large CD172^+^, mApple^+ve^ macrophages (**Figure S2A**). In WT and heterozygote recipients at 1 week, mApple^+^ donor cells were detected in 5/9 recipients, albeit at lower frequency (**Figure S2A**). However, at later time points, mApple^+^ cells were detected in only 1/10 recipients (**Figure S2B**).

In the same cohort we determined whether mApple^+ve^ donor cells could be detected in the blood at the early time point. As noted above, in rat peripheral blood, the large majority of monocytes are non-classical (His48^low^/CD43^high^) and these cells are selectively lost in the *Csf1rko* [14]. After one week post BMT in *Csf1rko* recipients, the mApple^+ve^/CD45^+^ cells in blood varied from 0.01 to 1.5% (median 0.36%, n=7). By contrast, no mApple^+ve^ positive cells were detected in the blood in either WT or Het recipients of mApple^+ve^ bone marrow (total n=9) (**Figure S3B-D**). The majority of the mApple^+ve^ cells detected in the blood of the *Csf1rko* recipients resembled the non-classical monocytes (His48^low^/CD43^high^) whereas a greater proportion of residual recipient monocytes were His48^hi^/Cd43^low^ (**Figure S2D**). In summary, both sustained population of the peritoneal cavity and transient donor cell egress were detectable only in the *Csf1rko*. The proliferation and differentiation of donor cells in the *Csf1rko* peritoneal cavity is presumably driven by CSF1. Cultivation of rat bone marrow cells in CSF1 for 7 days *in vitro* generate pure cultures of proliferating bone marrow-derived macrophages (BMDM)[29]. We considered the possibility that cultivation of bone marrow in CSF1 prior to transfer would accelerate phenotypic rescue. However, transfer of up to 10 × 10^6^ BMDM failed to achieve engraftment or survival in 8/8 of *Csf1rko* recipients.

Turley *et al* [30] reported that spleen cells injected into the mouse peritoneal cavity preferentially access the pancreatic lymph nodes, whilst Jackson-Jones *et al* [31] emphasised uptake into fat-associated lymphoid clusters in the omentum. In *Csf1rko* rat recipients, large foci of invading mApple^+ve^ cells could be detected by whole mount imaging of tissues throughout the peritoneal cavity. **Figure 5A** shows the clusters of mApple^+ve^ cells in presumptive milky spots in the greater omentum (5A(i)) as well as the stellate macrophage-like cells in the intervening omental membrane (5A(ii)). In other areas, the pattern strongly suggested focal invasion of small numbers of donor cells followed by local proliferation and migration to occupy the vacant macrophage niche. For example, **Figure 5B** shows early stages of repopulation of the diaphragm. mApple^+ve^ cells appear to have extravasated from a vessel and migrated into the tissue to occupy typical locations of macrophages aligned with the muscle fibres. Similarly, in **Figure 5C**, several lobules of the epididymal fat pad contain mApple^+ve^ cells that have adopted a typical distribution of adipose tissue macrophages whilst adjacent adipose lobules remain macrophage-deficient. Taken together these images are consistent with multifocal invasion of donor cells through the mesothelial lining. In tissues outside of the peritoneal cavity at this time point we detected occasional foci of donor cells that are also strongly suggestive of seeding and ongoing local proliferation. **Figure 5D** shows an example in the lung. No mApple^+^ foci were detected in the peritoneal cavity or tissues analysed from either heterozygous or WT marrow recipients indicating that donor cells require a vacant niche in order to engraft (data not shown).

**Figure 5:**
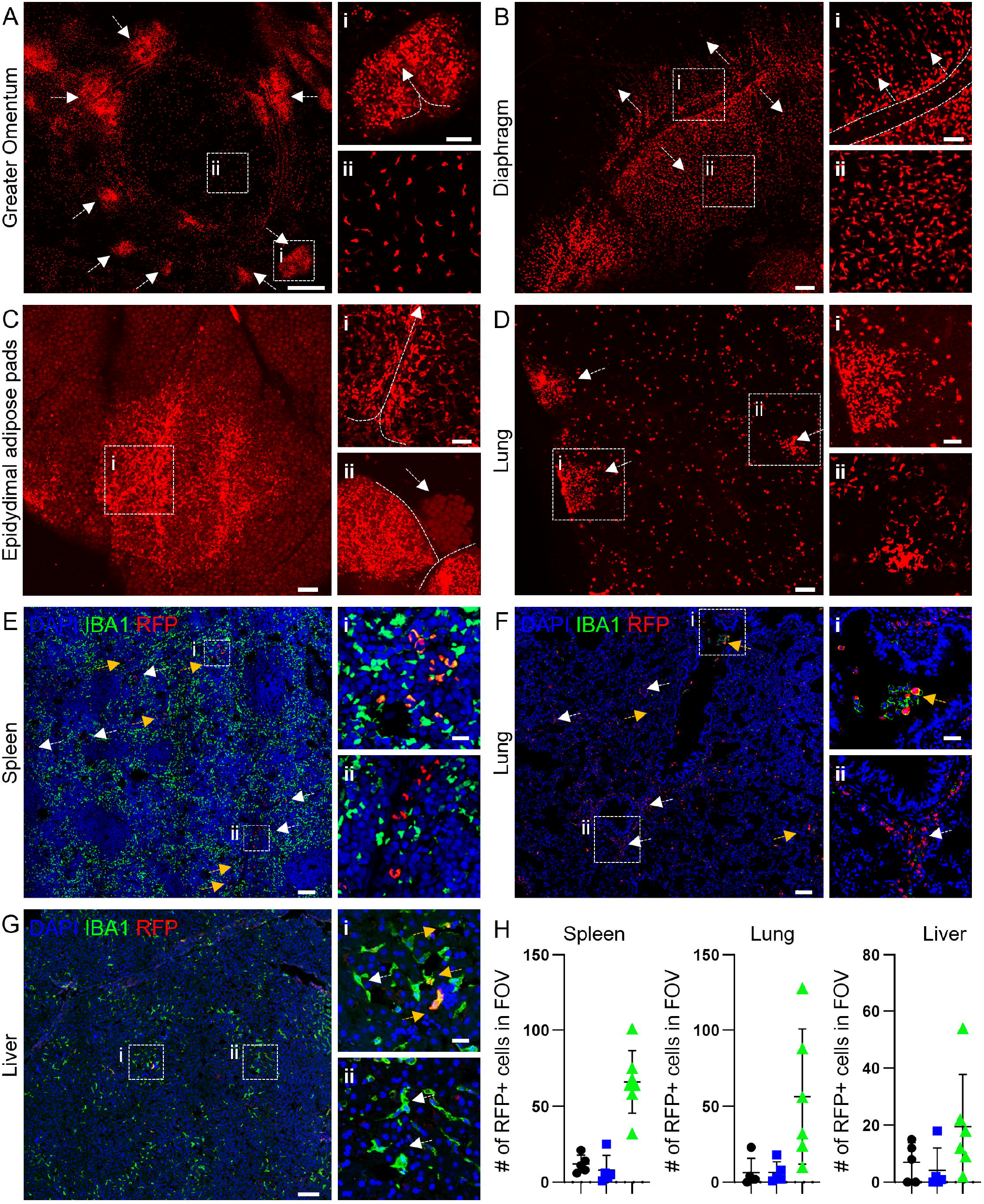
Spatial analysis of donor cells invasion 1-week post-transplant. *Csf1r*^+/+^ (WT) and *Csf1r*^+/-^ (Het) and *Csf1rko* rats received 2×10^7^ WT *Csf1r*^*mApple*^ female bone marrow (BM) cells by intraperitoneal injection at 2 weeks of age and were harvested 1-week later (A–D) Representative whole mounted tissues showing examples of the pattern of infiltration and aggregation of *Csf1r*^mApple^ donor cells in *Csf1rko* recipient rats one week post BMT (A) Greater omentum showing infiltration of donor cells in milky spots (i) and spatially separated tissue resident macrophages in the omental membrane (ii). (B) Diaphragm showing infiltration of donor cells through vessel structures (i) and spatially separated tissue resident macrophages interspersed between fibres (ii). (C) Epidydimal adipose fat pad showing infiltration into adipose tissue through a vessel structure (i). In the inset (ii) the white arrow highlights a lobule detectable with faint autofluorescence that contains no infiltrating mApple^+ve^ cells, adjacent to two fully repopulated lobules. (D) Foci of infiltrating donor cells on the edge/surface of the lungs (i) and within the lung parenchyma (ii). Note also individual positive cells throughout the parenchyma. (E–G) Immunolabelling of IBA1 (green), red fluorescent protein (RFP; mApple) and DAPI (blue) showing infiltration of donor *Csf1r*^mApple^ cells in (E) spleen, (F) lung and (G) liver. In all tissues, foci of IBA1^+^RFP^+^ (i; orange arrows) and IBA1^-^/RFP^+^ (ii; white arrows) cells are detected. (H) Quantification of total IBA1^-^/RFP^+^ cells in field of view (FOV) in spleen, lung, and liver of donor *Csf1r*^mApple^ infiltration in *Csf1r*^+/+^ (WT; black circles) *Csf1r*^+/-^ (Het; blue squares), *Csf1rko* (green triangles). Each data point represents mean of number of IBA1^-^/RFP^+^ cells from 4 different fields of view from at least 2 sectional depths per animal, n = 5-6 per group. Scale bars: A) 500μm, inset = 100μm; B-D) 200μm, inset = 100μm; E-G) 100μm, inset = 20μm.

*Aif1* mRNA (encoding IBA1) is not expressed by myeloid progenitor cells (see for example, data in BioGPS.org) but is induced during differentiation in response to CSF1 and is highly-expressed by rat BMDM generated *in vitro* [29]. The recent generation of an *Aif1* knock-in reporter transgene in rats [32] confirmed that IBA1 is expressed by monocytes. Since the *Csf1r-*mApple transgene is expressed earlier in myeloid commitment [25] we hypothesised that double-labelling for IBA1 and mApple would give insight into the maturation state of cells repopulating tissue MPS populations. **Figure 5E-G** show clusters of infiltrating cells in spleen, lung and liver week post BMT. A consistent feature is the presence within the destination tissue of mApple^+ve^ cells that lack detectable IBA1, implying that they are not derived from IBA1^+^ monocytes. These cells are most obvious in the lung (**Figure 5F**) where there are fewer residual IBA1^+^ cells in the *Csf1rko* recipient. The mApple^+ve^/IBA1^-ve^ cells were detected in spleen, lung and liver only in the *Csf1rko* BMT rats (**Figure 5H**) further illustrating the lack of engraftment in WT and heterozygous recipients in which the niche is occupied.

## Discussion

The generation and maintenance of cells of the mononuclear phagocyte system depends on signals from CSF1R in response to its ligands, CSF1 and IL34. CSF1 as the sole stimulus promotes the proliferation and differentiation of rat macrophages *in vitro* [29]. Because the ligands are internalised and degraded by CSF1R-mediated endocytosis, there is an intrinsic homeostatic mechanism by which macrophages control the availability of their own growth factors In the *Csf1rko* rat, tissue and circulating levels of CSF1 in the circulation increase massively and are progressively reduced to basal levels as macrophage populations are restored [14].

There have been numerous published studies of the ability of bone marrow-derived cells to repopulate vacant niches in individual organs in the mouse [1, 3]. The current study demonstrates that the vacant macrophage niche in every organ of the *Csf1rko* rat can be repopulated by bone marrow-derived progenitors. Wild-type bone marrow cells transferred into the peritoneum of the *Csf1rko* rat rapidly proliferate to repopulate typical large *Csf1r-*mApple^high^, CD172^+^, CD4^high^ peritoneal macrophages (**Figure S2**) and invade the mesentery, fat deposits and omental milky spots in the cavity and migrate to distant sites. Outside of the cavity, the donor cells migrated to achieve a regular distribution and appearance that closely resembles the density and distribution in the tissues of the *Csf1r*-mApple donors [25]. Our previous study showed that BMT does not restore CSF1-responsiveness in the bone marrow despite the excavation of the bone marrow cavity and restoration of bone marrow cellularity [14]. The lack of contribution of mApple^+^ donor cells to the hematopoietic stem cell compartment is further confirmed herein from the lack of expression of the reporter in neutrophils (which share a progenitor with monocytes [33]) and B cells as well as the absence from monocytes **(Figure 2)**. As a consequence of the lack of label in neutrophils and B cells it is possible to visualize the distribution of donor-derived cells in lymphoid organs of the recipients **(Figure 3)**, whereas the expression in B cells masks detection of the macrophage expression in the donor [25]. As well as restoring populations of macrophages that are absent in the *Csf1rko*, the mApple^+ve^ cells take over locations that were not entirely macrophage-deficient in the recipient, notably in the spleen, implying that the CSF1R-expressing wild-type cells have a proliferative advantage, likely because of high local CSF1 concentration in the red pulp [34]. That advantage includes repopulation of the entire lung macrophage population, both alveolar and interstitial, where the effect of the *Csf1rko* on macrophage numbers was least penetrant [14]. Lung interstitial and alveolar macrophage populations depend primarily upon granulocyte-macrophage colony stimulating factor (CSF2) [35]. However, CSF1R is highly expressed in lung macrophages in rats and mice [4, 29] and respiratory distress is the major welfare concern requiring euthanasia in *Csf1rko* animals.

One location where cells that were mApple^+ve^ in the donor remained mApple^-ve^ in the rescued *Csf1rko* rats was in the T cell areas of spleen (PALS) (**Figure 4**). IBA1^+^ cells in Peyer’s patches and in white pulp of lymph nodes were also not depleted in the *Csf1rko* [14]. These cells are likely splenic classical dendritic cells which express the *Csf1r-*mApple transgene but probably depend upon FLT3, rather than CSF1R [36]. Classical dendritic cells in the rat have not been well-defined [2] and there are currently no definitive markers. However, FLT3 ligand (*Flt3lg*) is highly expressed by rat T cells ([37]; biogps.org/ratatlas) so it is likely that these CSF1R-independent populations are maintained by FLT3 signalling.

The trafficking of donor cells in this model has some parallels with tumor metastasis. The omentum is a major site of initial metastasis of ovarian cancer and invasion is supported by omental macrophages [38, 39]. In response to inflammation in the peritoneal cavity, resident peritoneal macrophages can migrate from the cavity to the milky spots on the omentum, a visceral adipose tissue [40, 41]. Based upon the analysis of visceral tissues one week post-BMT, this appears to be one route of trafficking of at least some donor-derived progenitor cells. However, invasion of visceral fat and the diaphragm were also evident at this timepoint **(Figure 5)**, and direct trafficking to the pancreatic lymph nodes has also been reported in mice [30]. From these initial locations, we hypothesise that donor cells migrate through the lymphatics to enter the blood and then seed other organs.

The rescue of the *Csf1rko* rat by BMT without conditioning has some parallels with similar studies in the mouse [22]. A major difference is that in the mouse, BMT was carried out in neonates, before any overt developmental defects. By contrast, in the rat BMT was carried out at weaning, when the severe effects of the *Csf1rko* on growth and maturation of organs are already evident [14]. The focus in the mouse studies was on repopulation of microglia in the brain. Interestingly, the authors showed that repopulation of the brain was independent of CCR2, but nevertheless could be achieved by transfer of purified blood monocytes stringently depleted of progenitors. They did not address the question of whether donor cells contributed to sustained monocytopoiesis or repopulation of other tissue macrophage populations.

In the rat model, the pattern of migration observed at the early time points and the cellular phenotype of early foci (IBA1^-ve^/mApple^+ve^, **Figure 5E-H**) suggests that each organ is seeded by a small number of relatively immature cells which then expand, differentiate and migrate in response to locally-produced CSF1 [9] until a local balance between CSF1 production and consumption is achieved. Small numbers of mApple^+ve^ myeloid cells were detected in peripheral blood at one week post BMT (**Figure S3**). Based upon the surface markers (CD172^+^/His48^low^/ CD43^high^) on these cells, they are most likely non-classical monocytes derived from the donor marrow, but that population is clearly not sustained from donor origin in the long term BMT recipients (**Figure 2**). Although they could contribute, we consider it unlikely that these monocyte-like mApple^+ve^ cells are the main source of repopulating cells in destination organs. The experiment in **Figure S2** indicates that donor cells proliferate rapidly to populate the peritoneal cavity in the *Csf1rko* and the lack of sustained donor contribution in the WT and Het recipients indicates that proliferation requires a vacant niche. Niche competition was previously modelled *in vitro*, in studies showing that small numbers of peritoneal macrophages can compete for CSF1 and suppress proliferation of BMDM [42]

The lack of IBA1 expression in infiltrating mApple^+ve^ cells (**Figure 5**) and the pattern of repopulation from local foci suggests that the migratory cells that populate every organ are immature and have a high proliferative potential; facilitated by the elevated levels of CSF1 in *Csf1rko* recipients [14]. Since the only marker we have, *Csf1r-*mApple, is itself induced during macrophage lineage commitment and differentiation [25], we cannot eliminate the possibility that the migratory cells lack the reporter and acquire it in the destination organ. Furthermore, CSF1/IL34 may not be the only signals promoting migration and tissue infiltration in the *Csf1rko*. The granulocytosis we observe suggests an increase in availability of CSF3 (G-CSF), which is commonly used to mobilise stem cells [43]. One possibility is that committed progenitor cells in the transferred marrow migrate directly to distal sites. The presence of macrophage colony-forming precursors with high proliferative potential in bone marrow was recognised previously in mice [44]. Their ability to mobilise through the blood is evident from the identification of macrophage colony forming cells in peritoneal inflammatory peritoneal exudates [45]. In the chick, a progenitor in bone marrow can generate macrophage-restricted chimerism in adult birds when injected *in ovo* [6]. The progenitors may be related to the committed monocyte progenitor described in mouse bone marrow by Hettinger *et al*. [46], which could contribute to monocyte and tissue macrophage populations upon adoptive transfer.

The prevailing view of the homeostatic maintenance of resident tissue macrophages based upon lineage trace models in inbred mice is that most populations are seeded during embryonic development and maintained by self-renewal without substantial input from bone marrow-derived blood monocytes [1, 3, 47]. Once established in the rescued *Csf1rko* the donor-derived resident macrophages in every organ must be sustained by self-renewal since there are no mApple^+ve^ monocytes. However, this likely depends in part upon increased availability of CSF1 from the circulation in the absence of consumption by blood monocytes [9, 48]. The *Csf1rko* rat rescued by bone marrow transfer remains deficient in long-lived non-classical or patrolling monocyte population that is proposed to monitor endothelial integrity [49, 50]. Neither monocytes nor bone marrow progenitors in these rescued *Csf1rko* animals express CSF1R. Yet, these rescued rats are long-lived and fertile. Accordingly, they provide a unique model to understand monocyte-macrophage homeostasis.

## Materials and Methods

### Ethics Statement

Rats were bred and maintained in specific pathogen free facilities at The University of Queensland (UQ) under protocols approved by the UQ Animal Ethics Unit (Approval MRI-UQ/424/18, MRI-UQ/318/17).

### Generation of transgenic rats and animal husbandry

To generate animals in which we could transfer *Csf1r-*mApple bone marrow to *Csf1rko* recipients we crossed *Csf1r*^*+/-*^ and *Csf1r*^mApple^ rats, both on the SD background. We chose a male and female founder carrying both the knockout allele and reporter, and brother-sister mated their progeny with selection for both markers for a further 3 generations. We then separated the rats carrying only the *Csf1r*^mApple^ transgene to act as donors and used heterozygous mating of their siblings to generate *Csf1rko* recipients. The resultant inbred *Csf1rko*^SD^ rats did not develop teeth and to ensure their survival and maximise their growth, we established the same maintenance approach as previously used in the *Csf1rko*^DA^ line [14] and commenced a feeding regime including wet mashed standard chow and a veterinary powdered milk nutritional supplement (Di-Vetelact, Sydney, Australia). Littermates were kept in the same cage and had access to the same feeding supplements ad libitum. Animals were weighted daily with frequency reduced after 10 weeks to every 2 days.

### Bone marrow transfer (BMT)

Bone marrow was obtained from the femurs and tibias of *Csf1r* wild-type transgenic *Csf1r-*mApple (*Csf1r*^mApple^) reporter rats [25] by flushing with a 26G needle. 2×10^7^ bone marrow cells in 0.9% sodium chloride solution supplemented with 2% fetal bovine serum (FBS) were transferred to 2-week-old *Csf1rko*^SD^ recipients by intraperitoneal injection. Recipients and littermates (non-transferred) were maintained on the supplemented diet. To investigate temporal dynamics of donor cell infiltration, 2×10^7^ cells were also transferred into WT or heterozygote (*Csf1r*^*+/-*^) littermates which were harvested for investigation one week post-BMT.

### IGF1 enzyme-linked immunosorbent assay

Serum IGF1 was measured using a Mouse/Rat DuoSet ELISA kit (R&D Systems, Minneapolis, Minnesota, USA) according to manufacturer’s instructions as previously described [14]. Colorimetric changes were read using an automated microplate reader with Gen5 Microplate Reader Software (Synergy H1 Hybrid Reader, BioTek, Vermont, USA).

### Immunohistochemistry (IHC) and Immunofluorescence (IF) microscopy

Tissues for IHC were harvested and fixed in 4% paraformaldehyde for 24 hours and then processed for paraffin-embedded histology using routine methods. Sections were deparaffinised and rehydrated in descending ethanol series. Tissues for IF were fixed in 4% paraformaldehyde for 2 hours and processed by sucrose saturation in 15-30% sucrose gradient over 2 days before embedding in Optimal Cutting Temperature compound (OCT, ProSciTech, Kirwan, Australia) and sectioning at 5 µm thickness on Lieca 1950 cryostat (Mount Waverley, Australia). Ki67, IBA1 and RFP epitope retrieval was performed by heat induction in Diva Decloaker (DV2004MX, Lot:011519, Biocare Medical, California, USA). Sections were stained with rabbit anti-Ki67 (Abcam ab16667, Cambridge, UK) or rabbit anti-IBA1 (FUJIFILM 019–19741, Wako Chemicals Richmond, Virginia, USA), or rabbit anti-RFP (Abcam ab124754, Cambridge, UK). For IHC, secondary detection was with DAKO Envision anti-rabbit HRP detection reagents (Agilent Technologies Australia, Mulgrave, Australia). Sections were counterstained hematoxylin (Sigma-Aldrich, St. Louis, USA) for 1 minute. Sections were then dehydrated in ascending ethanol series, clarified with xylene and mounted with DPX mountant (Sigma-Aldrich, St. Louis, USA). Whole-slide digital imaging of IHC sections was performed on the VS120 Olympus slide scanner (Olympus Corporation, Tokyo, Japan). For IF, unconjugated primary antibodies were detected with fluorescently-labelled species appropriate secondary antibodies including Goat Anti-Rabbit IgG (H+L) Alexa Fluor 488 and Goat Anti-Rat IgG (H+L) Alexa Fluor 647 (ThermoFisher Scientific, Brisbane, Australia). All IF sections were counterstained with 4′,6-diamidino-2-phenylindole (DAPI; Thermo Fisher Scientific, USA). Immunofluorescent images were acquired on an Olympus FV3000 microscope. For wholemount imaging, tissues were left as undisturbed as possible for direct imaging through the tissue surface. For tissues with reduced laser penetration (such as spleen or kidney) tissues were trimmed to a thickness of 2 mm and imaged inwards from the capsular surface. *Csf1r-*mApple signal was acquired using the 561-diode laser on the FV3000 main combiner (FV31-MCOMB). For DAPI, AF488 or AF647, signal was acquired using the 405-, 488-and 633-diode lasers respectively on the FV3000 main combiner (FV31-MCOMB). Morphometric analysis of either DAB-positive areas or IF secondary antibody staining was quantified using ImageJ (https://imagej.net/) [51]. Colocalization analysis for IBA1 and RFP was performed using the Coloc 2 ImageJ plugin [52]. All images were coded and assessed blindly at least 3 sectional depths. Background intensity thresholds were applied using an ImageJ macro which measures pixel intensity across all immunostained and non-stained areas of the images and these were converted to percent staining area.

### Flow cytometry

100 μl peripheral blood was collected into ethylenediaminetetraacetic acid (EDTA) tubes by cardiac puncture after carbon dioxide euthanasia. Red cells were lysed for 2 min in ACK lysis buffer (150mM NH_4_Cl, 10mM KHCO_3_, 0.1mM EDTA, pH7.4), the leukocytes were centrifuged and washed twice in PBS and the pellet resuspended in flow cytometry (FC) buffer (PBS/2% FBS) for staining. Peritoneal flushes were collected in PBS, centrifuged, and resuspended in FC buffer for staining. BM cells were isolated by crushing hind limbs using a mortar and pestle and processed for staining in FC buffer. Cells were stained for 45 min on ice in FC buffer containing unlabelled CD32 (BD Biosciences, Sydney, Australia) to block Fc receptor binding, HIS48-FITC, CD11B/C-BV510, CD45R-BV786, CD45-PE/Cy7 (BD Biosciences), CD43-AF647, CD4-APC/Cy7, (Biolegend, San Diego, CA, USA) and CD172A-AF405 (Novus, Noble Park North, Australia). Cells were washed twice and resuspended in FC buffer containing 7AAD (LifeTechnologies, Musgrave, Australia) for acquisition on either a Cytoflex (Beckman Coulter Life Sciences, Sydney, Australia) or Fortessa (Becton Dickinson) flow cytometer. Relevant single-color controls were used for compensation and unstained and fluorescence-minus-one controls were used to confirm gating strategies. Flow cytometry data were analyzed using FlowJo 10 (Tree Star). Live single cells were identified for phenotypic analysis by excluding doublets (FSC-A > FSC-H), 7AAD^+^ dead cells and debris.

### Statistical Analysis

Graphing and statistical analyses were conducted using GraphPad Prism 8.3.1 (www.graphpad.com). All sample sizes and statistical tests employed are documented in Figure legends.

## Legends to Figures

**Supplementary Figure 1. Flow cytometry gating strategy**.

Bone marrow, peripheral blood and peritoneal cells were harvested from donor *Csf1r*^mApple^ rats and stained for the indicated markers as described in Materials and Methods.

**Supplementary Figure 2. Repopulation of the peritoneal cavity with donor *Csf1r***^**mApple**^ **cells in WT, Hetero*Csf1rko* rats after BMT**.

*Csf1r*^+/+^ (WT) and *Csf1r*^+/-^ (Het) and *Csf1rko* rats were transplanted with 2×10^7^ WT *Csf1r*^mApple^ bone marrow cells intraperitoneally (BMT) at 2 weeks of age and peritoneal cells were harvested by lavage at 1 week (Panel A) or 2 or 4 weeks post-transplant (Panel B). Within one week in the *Csf1rko* peritoneal CD172^+ve^ macrophages, which are absent in these animals, were repopulated entirely from donor origin. In half of the WT/HET recipients, there were no detectable mApple^+^ cells (typical profile shown). In the remainder, there was a significant donor population (CD45^+^/mApple^+^). as shown in the histogram. In an additional cohort of WT and HET recipients assayed at 2-4 weeks (data combined in Panel B), donor mApple^+^ cells were no longer detected with a single exception at 4 weeks in which 25% of cells had low but detectable expression. The profile of peritoneal cells in this recipient is shown at right.

**Supplementary Figure 3. Circulating *Csf1r***^**mApple**^ **cells in *Csf1rko* rats one week post BMT**. *Csf1r*^+/+^ (WT) and *Csf1r*^+/-^ (Het) (combined n=9) and *Csf1rko* (n=7) rats were transplanted with 2×10^7^ WT *Csf1r*^mApple^ bone marrow cells intraperitoneally (BMT) at 2 weeks of age and harvested 1-week post-transplant. (A) mApple^+^ cell profile in WT donor bone marrow (BM), comprising ∼13% CD172a^Low^/HIS48^+^ neutrophils, 12% CD172^+^/CD43^+^/HIS48^+^ monocytes and ∼84% CD45R+ B cells. (B) mApple^+^ cells in peripheral blood (as % of Live CD45^+^ cells). Lines indicate median and upper/lower quartiles. (C) Profile of mApple^-ve^ leukocytes in peripheral blood of a representative *Csf1r*^+/-^ (Het) recipient. (D) Profile of mApple^+ve^ and mApple^-ve^ leukocytes in peripheral blood of a representative *Csf1rko* recipient.

## Acknowledgements

DAH and KMI are grateful for core laboratory support from the Mater Foundation. We appreciate the support of the Preclinical Imaging, Biological Resources, Histology, Microscopy and Flow Cytometry Core Facilities at the Translational Research Institute. We would like to thank Ms. Lisa Foster (Manager, UQ PACE Biological Resources Facility) and her staff (especially Rachel Smith) for excellent support with breeding and husbandry of the *Csf1rko* rats. This work was funded by an Australian Research Council Grant DP20013245 to DAH and KMI and an NHMRC Investigator Grant 2009750 to DAH.

## Author contributions

Conceptualisation: DAH, KMI. Investigation: AS, DCC, SK, OP, SH Writing and editing AS, KMS, DAH, KMI. Supervision: KMS, DAH, KMI. Funding Acquisition: DAH, KMI.

## References

1 Guilliams, M., Thierry, G. R., Bonnardel, J. and Bajenoff, M., Establishment and Maintenance of the Macrophage Niche. Immunity 2020. 52: 434–451.

2 Hume, D. A., Caruso, M., Keshvari, S., Patkar, O. L., Sehgal, A., Bush, S. J., Summers, K. M., Pridans, C. and Irvine, K. M., The mononuclear phagocyte system of the rat. J. Immunol 2021. 206: 2251–2263.

3 Hume, D. A., Irvine, K. M. and Pridans, C., The Mononuclear Phagocyte System: The Relationship between Monocytes and Macrophages. Trends Immunol 2018. 40: 98–112.

4 Summers, K. M., Bush, S. J. and Hume, D. A., Transcriptional network analysis of transcriptomic diversity in resident tissue macrophages and dendritic cells in the mouse mononuclear phagocyte system. PloS.Biol. 2020. 18: e3000859.

5 Chitu, V. and Stanley, E. R., Regulation of Embryonic and Postnatal Development by the CSF-1 Receptor. Curr Top Dev Biol 2017. 123: 229–275.

6 Garceau, V., Balic, A., Garcia-Morales, C., Sauter, K. A., McGrew, M. J., Smith, J., Vervelde, L., Sherman, A., Fuller, T. E., Oliphant, T., Shelley, J. A., Tiwari, R., Wilson, T. L., Chintoan-Uta, C., Burt, D. W., Stevens, M. P., Sang, H. M. and Hume, D. A., The development and maintenance of the mononuclear phagocyte system of the chick is controlled by signals from the macrophage colony-stimulating factor receptor. BMC Biol 2015. 13: 12.

7 Garceau, V., Smith, J., Paton, I. R., Davey, M., Fares, M. A., Sester, D. P., Burt, D. W. and Hume, D. A., Pivotal Advance: Avian colony-stimulating factor 1 (CSF-1), interleukin-34 (IL-34), and CSF-1 receptor genes and gene products. J Leukoc Biol 2010. 87: 753–764.

8 Hume, D. A., Caruso, M., Ferrari-Cestari, M., Summers, K. M., Pridans, C. and Irvine, K. M., Phenotypic impacts of CSF1R deficiencies in humans and model organisms. J Leukoc Biol 2020. 107: 205–219.

9 Sehgal, A., Irvine, K. M. and Hume, D. A., Functions of macrophage colony-stimulating factor (CSF1) in development, homeostasis, and tissue repair. Semin Immunol 2021. 54: 101509.

10 Lichanska, A. M., Browne, C. M., Henkel, G. W., Murphy, K. M., Ostrowski, M. C., McKercher, S. R., Maki, R. A. and Hume, D. A., Differentiation of the mononuclear phagocyte system during mouse embryogenesis: the role of transcription factor PU.1. Blood 1999. 94: 127–138.

11 Henson, P. M. and Hume, D. A., Apoptotic cell removal in development and tissue homeostasis. Trends Immunol 2006. 27: 244–250.

12 Munro, D. A. D., Bradford, B. M., Mariani, S. A., Hampton, D. W., Vink, C. S., Chandran, S., Hume, D. A., Pridans, C. and Priller, J., CNS macrophages differentially rely on an intronic Csf1r enhancer for their development. Development 2020. 147: dev194449.

13 Rojo, R., Raper, A., Ozdemir, D. D., Lefevre, L., Grabert, K., Wollscheid-Lengeling, E., Bradford, B., Caruso, M., Gazova, I., Sanchez, A., Lisowski, Z. M., Alves, J., Molina-Gonzalez, I., Davtyan, H., Lodge, R. J., Glover, J. D., Wallace, R., Munro, D. A. D., David, E., Amit, I., Miron, V. E., Priller, J., Jenkins, S. J., Hardingham, G. E., Blurton-Jones, M., Mabbott, N. A., Summers, K. M., Hohenstein, P., Hume, D. A. and Pridans, C., Deletion of a Csf1r enhancer selectively impacts CSF1R expression and development of tissue macrophage populations. Nat Commun 2019. 10: 3215.

14 Keshvari, S., Caruso, M., Teakle, N., Batoon, L., Sehgal, A., Patkar, O. L., Ferrari-Cestari, M., Snell, C. E., Chen, C., Stevenson, A., Davis, F. M., Bush, S. J., Pridans, C., Summers, K. M., Pettit, A. R., Irvine, K. M. and Hume, D. A., CSF1R-dependent macrophages control postnatal somatic growth and organ maturation. PLoS Genet 2021. 17: e1009605.

15 Erblich, B., Zhu, L., Etgen, A. M., Dobrenis, K. and Pollard, J. W., Absence of colony stimulation factor-1 receptor results in loss of microglia, disrupted brain development and olfactory deficits. PLoS One 2011. 6: e26317.

16 Pridans, C., Raper, A., David, G. M., Alves, J., Sauter, K. A., Lefevre, L., Regan, T., Grabert, K., Meek, S., Sutherland, L., Thomson, A. J., Clohisey, S. M., Rojo, R., Lisowski, Z. M., Wallace, R., Upton, K. R., Tsai, Y. T., Brown, D., Smith, L. B., Mabbott, N. A., Picardo, P., Cheeseman, M. T., Burdon, T. and Hume, D. A., Pleiotropic Impacts of Macrophage and Microglial Deficiency on Development in Rats with Targeted Mutation of the Csf1r Locus. J.Immunol. 2018. 201(9): 2683–2699.

17 Patkar, O. L., Caruso, M., Teakle, N., Keshvari, S., Bush, S. J., Pridans, C., Belmer, A., Summers, K. M., Irvine, K. M. and Hume, D. A., Analysis of homozygous and heterozygous Csf1r knockout in the rat as a model for understanding microglial function in brain development and the impacts of human CSF1R mutations. Neurobiol Dis 2021: 105268.

18 Guo, L., Bertola, D. R., Takanohashi, A., Saito, A., Segawa, Y., Yokota, T., Ishibashi, S., Nishida, Y., Yamamoto, G. L., Franco, J., Honjo, R. S., Kim, C. A., Musso, C. M., Timmons, M., Pizzino, A., Taft, R. J., Lajoie, B., Knight, M. A., Fischbeck, K. H., Singleton, A. B., Ferreira, C. R., Wang, Z., Yan, L., Garbern, J. Y., Simsek-Kiper, P. O., Ohashi, H., Robey, P. G., Boyde, A., Matsumoto, N., Miyake, N., Spranger, J., Schiffmann, R., Vanderver, A., Nishimura, G., Passos-Bueno, M., Simons, C., Ishikawa, K. and Ikegawa, S., Bi-allelic CSF1R Mutations Cause Skeletal Dysplasia of Dysosteosclerosis-Pyle Disease Spectrum and Degenerative Encephalopathy with Brain Malformation. Am J Hum Genet 2019. 104: 925–935.

19 Oosterhof, N., Chang, I. J., Karimiani, E. G., Kuil, L. E., Jensen, D. M., Daza, R., Young, E., Astle, L., van der Linde, H. C., Shivaram, G. M., Demmers, J., Latimer, C. S., Keene, C. D., Loter, E., Maroofian, R., van Ham, T. J., Hevner, R. F. and Bennett, J. T., Homozygous Mutations in CSF1R Cause a Pediatric-Onset Leukoencephalopathy and Can Result in Congenital Absence of Microglia. Am J Hum Genet 2019. 104: 936–947.

20 Kindis, E., Simsek-Kiper, P. O., Kosukcu, C., Taskiran, E. Z., Gocmen, R., Utine, E., Haliloglu, G., Boduroglu, K. and Alikasifoglu, M., Further expanding the mutational spectrum of brain abnormalities, neurodegeneration, and dysosteosclerosis: A rare disorder with neurologic regression and skeletal features. Am J Med Genet A 2021. 185: 1888–1896.

21 Tamhankar, P. M., Zhu, B., Tamhankar, V. P., Mithbawkar, S., Seabra, L., Livingston, J. H., Ikeuchi, T. and Crow, Y. J., A Novel Hypomorphic CSF1R Gene Mutation in the Biallelic State Leading to Fatal Childhood Neurodegeneration. Neuropediatrics 2020. 51: 302–306.

22 Bennett, F. C., Bennett, M. L., Yaqoob, F., Mulinyawe, S. B., Grant, G. A., Hayden Gephart, M., Plowey, E. D. and Barres, B. A., A Combination of Ontogeny and CNS Environment Establishes Microglial Identity. Neuron 2018. 98: 1170–1183 e1178.

23 Beers, D. R., Henkel, J. S., Xiao, Q., Zhao, W., Wang, J., Yen, A. A., Siklos, L., McKercher, S. R. and Appel, S. H., Wild-type microglia extend survival in PU.1 knockout mice with familial amyotrophic lateral sclerosis. Proc Natl Acad Sci U S A 2006. 103: 16021–16026.

24 MacDonald, K. P., Palmer, J. S., Cronau, S., Seppanen, E., Olver, S., Raffelt, N. C., Kuns, R., Pettit, A. R., Clouston, A., Wainwright, B., Branstetter, D., Smith, J., Paxton, R. J., Cerretti, D. P., Bonham, L., Hill, G. R. and Hume, D. A., An antibody against the colony-stimulating factor 1 receptor depletes the resident subset of monocytes and tissue-and tumor-associated macrophages but does not inhibit inflammation. Blood 2010. 116: 3955–3963.

25 Irvine, K. M., Caruso, M., Cestari, M. F., Davis, G. M., Keshvari, S., Sehgal, A., Pridans, C. and Hume, D. A., Analysis of the impact of CSF-1 administration in adult rats using a novel Csf1r-mApple reporter gene. J Leukoc Biol 2020. 107: 221–235.

26 Grabert, K., Sehgal, A., Irvine, K. M., Wollscheid-Lengeling, E., Ozdemir, D. D., Stables, J., Luke, G. L., Ryan, M. D., Adamson, A., Humphreys, N. E., Sandrock, C. J., Verkasalo, V. A., Rojo, R., Mueller, W., Hohenstein, P., Pettit, A. L., Pridans, C. and Hume, D. A., A mouse transgenic line that reports CSF1R protein expression provides a definitive differentiation marker for the mouse mononuclear phagocyte system. J.Immunol. 2020. 205: 3154–3166.

27 Hawley, C. A., Rojo, R., Raper, A., Sauter, K. A., Lisowski, Z. M., Grabert, K., Bain, C. C., Davis, G. M., Louwe, P. A., Ostrowski, M. C., Hume, D. A., Pridans, C. and Jenkins, S. J., Csf1r-mApple Transgene Expression and Ligand Binding In Vivo Reveal Dynamics of CSF1R Expression within the Mononuclear Phagocyte System. J Immunol 2018. 200: 2209–2223.

28 Millard, S. M., Heng, O., Opperman, K. S., Sehgal, A., Irvine, K. M., Kaur, S., Sandrock, C. J., Wu, A. C., Magor, G. W., Batoon, L., Perkins, A. C., Noll, J. E., Zannettino, A. C. W., Sester, D. P., Levesque, J. P., Hume, D. A., Raggatt, L. J., Summers, K. M. and Pettit, A. R., Fragmentation of tissue-resident macrophages during isolation confounds analysis of single-cell preparations from mouse hematopoietic tissues. Cell Rep 2021. 37 110058.

29 Pridans, C., Irvine, K. M., Davis, G. M., Lefevre, L., Bush, S. J. and Hume, D. A., Transcriptomic Analysis of Rat Macrophages. Front Immunol 2020. 594594.

30 Turley, S. J., Lee, J. W., Dutton-Swain, N., Mathis, D. and Benoist, C., Endocrine self and gut non-self intersect in the pancreatic lymph nodes. Proc Natl Acad Sci U S A 2005. 102: 17729–17733.

31 Jackson-Jones, L. H., Smith, P., Portman, J. R., Magalhaes, M. S., Mylonas, K. J., Vermeren, M. M., Nixon, M., Henderson, B. E. P., Dobie, R., Vermeren, S., Denby, L., Henderson, N. C., Mole, D. J. and Benezech, C., Stromal Cells Covering Omental Fat-Associated Lymphoid Clusters Trigger Formation of Neutrophil Aggregates to Capture Peritoneal Contaminants. Immunity 2020. 52 700–715 e706.

32 VanRyzin, J. W., Arambula, S. E., Ashton, S. E., Blanchard, A. C., Burzinski, M. D., Davis, K. T., Edwards, S., Graham, E. L., Holley, A., Kight, K. E., Marquardt, A. E., Perez-Pouchoulen, M., Pickett, L. A., Reinl, E. L. and McCarthy, M. M., Generation of an Iba1-EGFP Transgenic Rat for the Study of Microglia in an Outbred Rodent Strain. eNeuro 2021. 8: ENEURO.0026-0021.2021.

33 Liu, Z., Gu, Y., Chakarov, S., Bleriot, C., Kwok, I., Chen, X., Shin, A., Huang, W., Dress, R. J., Dutertre, C. A., Schlitzer, A., Chen, J., Ng, L. G., Wang, H., Liu, Z., Su, B. and Ginhoux, F., Fate Mapping via Ms4a3-Expression History Traces Monocyte-Derived Cells. Cell 2019. 178: 1509–1525 e1519.

34 Bellomo, A., Mondor, I., Spinelli, L., Lagueyrie, M., Stewart, B. J., Brouilly, N., Malissen, B., Clatworthy, M. R. and Bajenoff, M., Reticular Fibroblasts Expressing the Transcription Factor WT1 Define a Stromal Niche that Maintains and Replenishes Splenic Red Pulp Macrophages. Immunity 2020. 53: 127–142 e127.

35 Shima, K., Arumugam, P., Sallese, A., Horio, Y., Ma, Y., Trapnell, C., Wessendarp, M., Chalk, C., McCarthy, C., Carey, B. C., Trapnell, B. C. and Suzuki, T., A murine model of hereditary pulmonary alveolar proteinosis caused by homozygous Csf2ra gene disruption. Am J Physiol Lung Cell Mol Physiol 2022. 322: L438–L448.

36 Durai, V., Bagadia, P., Briseno, C. G., Theisen, D. J., Iwata, A., Davidson, J. T. t., Gargaro, M., Fremont, D. H., Murphy, T. L. and Murphy, K. M., Altered compensatory cytokine signaling underlies the discrepancy between Flt3(-/-) and Flt3l(-/-) mice. J Exp Med 2018. 215: 1417–1435.

37 Summers, K. M., Bush, S. J., Wu, C. and Hume, D. A., Generation and network analysis of an RNA-seq transcriptional atlas for the rat. NAR Genom Bioinform 2022. 4: lqac017.

38 Etzerodt, A., Moulin, M., Doktor, T. K., Delfini, M., Mossadegh-Keller, N., Bajenoff, M., Sieweke, M. H., Moestrup, S. K., Auphan-Anezin, N. and Lawrence, T., Tissue-resident macrophages in omentum promote metastatic spread of ovarian cancer. J Exp Med 2020. 217: e20191869.

39 Krishnan, V., Tallapragada, S., Schaar, B., Kamat, K., Chanana, A. M., Zhang, Y., Patel, S., Parkash, V., Rinker-Schaeffer, C., Folkins, A. K., Rankin, E. B. and Dorigo, O., Omental macrophages secrete chemokine ligands that promote ovarian cancer colonization of the omentum via CCR1. Commun Biol 2020. 3: 524.

40 Meza-Perez, S. and Randall, T. D., Immunological Functions of the Omentum. Trends Immunol 2017. 38: 526–536.

41 Liu, M., Silva-Sanchez, A., Randall, T. D. and Meza-Perez, S., Specialized immune responses in the peritoneal cavity and omentum. J Leukoc Biol 2021. 109: 717–729.

42 Hume, D. A. and Gordon, S., The correlation between plasminogen activator activity and thymidine incorporation in mouse bone marrow-derived macrophages. Opposing actions of colony-stimulating factor, phorbol myristate acetate, dexamethasone and prostaglandin E. Exp Cell Res 1984. 150: 347–355.

43 Kaur, S., Sehgal, A., Wu, A. C., Millard, S. M., Batoon, L., Sandrock, C. J., Ferrari-Cestari, M., Levesque, J. P., Hume, D. A., Raggatt, L. J. and Pettit, A. R., Stable colony-stimulating factor 1 fusion protein treatment increases hematopoietic stem cell pool and enhances their mobilisation in mice. J Hematol Oncol 2021. 14: 3.

44 Kriegler, A. B., Verschoor, S. M., Bernardo, D. and Bertoncello, I., The relationship between different high proliferative potential colony-forming cells in mouse bone marrow. Exp Hematol 1994. 22: 432–440.

45 Chan, J., Leenen, P. J., Bertoncello, I., Nishikawa, S. I. and Hamilton, J. A., Macrophage lineage cells in inflammation: characterization by colony-stimulating factor-1 (CSF-1) receptor (c-Fms), ER-MP58, and ER-MP20 (Ly-6C) expression. Blood 1998. 92: 1423–1431.

46 Hettinger, J., Richards, D. M., Hansson, J., Barra, M. M., Joschko, A. C., Krijgsveld, J. and Feuerer, M., Origin of monocytes and macrophages in a committed progenitor. Nat Immunol 2013. 14: 821–830.

47 Ginhoux, F. and Guilliams, M., Tissue-Resident Macrophage Ontogeny and Homeostasis. Immunity 2016. 44: 439–449.

48 Yona, S., Kim, K. W., Wolf, Y., Mildner, A., Varol, D., Breker, M., Strauss-Ayali, D., Viukov, S., Guilliams, M., Misharin, A., Hume, D. A., Perlman, H., Malissen, B., Zelzer, E. and Jung, S., Fate mapping reveals origins and dynamics of monocytes and tissue macrophages under homeostasis. Immunity 2013. 38: 79–91.

49 Auffray, C., Fogg, D., Garfa, M., Elain, G., Join-Lambert, O., Kayal, S., Sarnacki, S., Cumano, A., Lauvau, G. and Geissmann, F., Monitoring of blood vessels and tissues by a population of monocytes with patrolling behavior. Science 2007. 317: 666–670.

50 Carlin, L. M., Stamatiades, E. G., Auffray, C., Hanna, R. N., Glover, L., Vizcay-Barrena, G., Hedrick, C. C., Cook, H. T., Diebold, S. and Geissmann, F., Nr4a1-dependent Ly6C(low) monocytes monitor endothelial cells and orchestrate their disposal. Cell 2013. 153: 362–375.

51 Inman, C. F., Rees, L. E., Barker, E., Haverson, K., Stokes, C. R. and Bailey, M., Validation of computer-assisted, pixel-based analysis of multiple-colour immunofluorescence histology. J Immunol Methods 2005. 302: 156–167.

52 Schindelin, J., Arganda-Carreras, I., Frise, E., Kaynig, V., Longair, M., Pietzsch, T., Preibisch, S., Rueden, C., Saalfeld, S., Schmid, B., Tinevez, J. Y., White, D. J., Hartenstein, V., Eliceiri, K., Tomancak, P. and Cardona, A., Fiji: an open-source platform for biological-image analysis. Nat Methods 2012. 9: 676–682.

