## Supplementary material for "Intraperitoneal transfer of wild-type bone marrow cells in the *Csf1r* knockout rat repopulates resident tissue macrophages without contributing to monocytopoiesis": Figures S1-S3

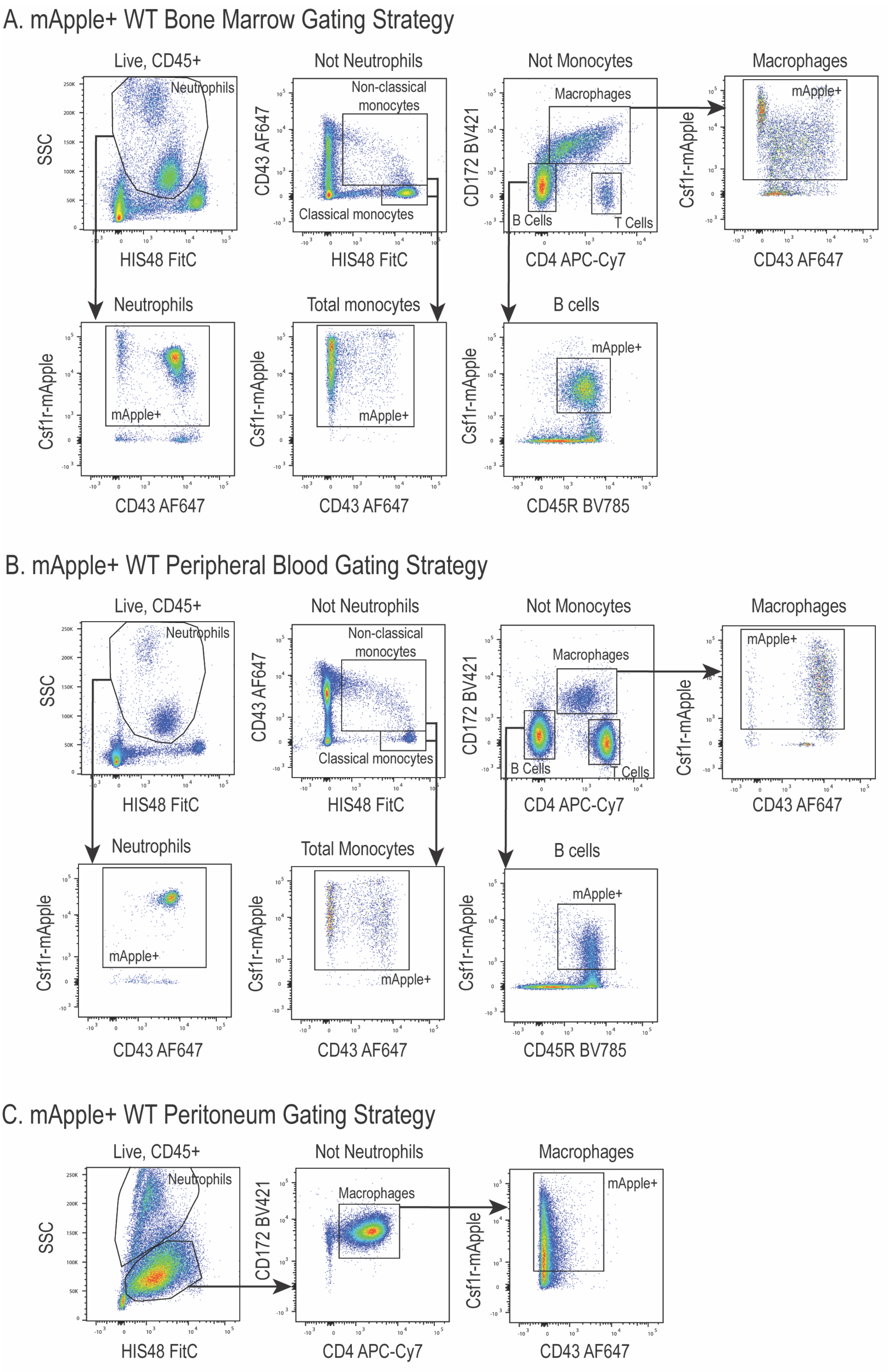
A. mApple+ WT Bone Marrow Gating Strategy
Live, CD45+
Neutrophils
SSC
HIS48 FitC
Not Neutrophils
Non-classical monocytes
Classical monocytes
CD43 AF647
Not Monocytes
Macrophages
B Cells
T Cells
CD172 BV421
CD4 APC-Cy7
Macrophages
mApple+
Csf1r-mApple
CD43 AF647
Neutrophils
mApple+
Total monocytes
B cells
CD45R BV785
B. mApple+ WT Peripheral Blood Gating Strategy
Live, CD45+
Neutrophils
SSC
HIS48 FitC
Not Neutrophils
Non-classical monocytes
Classical monocytes
CD43 AF647
Not Monocytes
Macrophages
B Cells
T Cells
CD172 BV421
CD4 APC-Cy7
Macrophages
mApple+
Csf1r-mApple
CD43 AF647
Neutrophils
Total Monocytes
B cells
CD45R BV785
C. mApple+ WT Peritoneum Gating Strategy
Live, CD45+
Neutrophils
SSC
HIS48 FitC
Not Neutrophils
Macrophages
CD172 BV421
CD4 APC-Cy7
Macrophages
mApple+
Csf1r-mApple
CD43 AF647

Figure S1

Figure S2

A. Peritoneal Csf1r-mApple+ cells 1 wk post-BMT

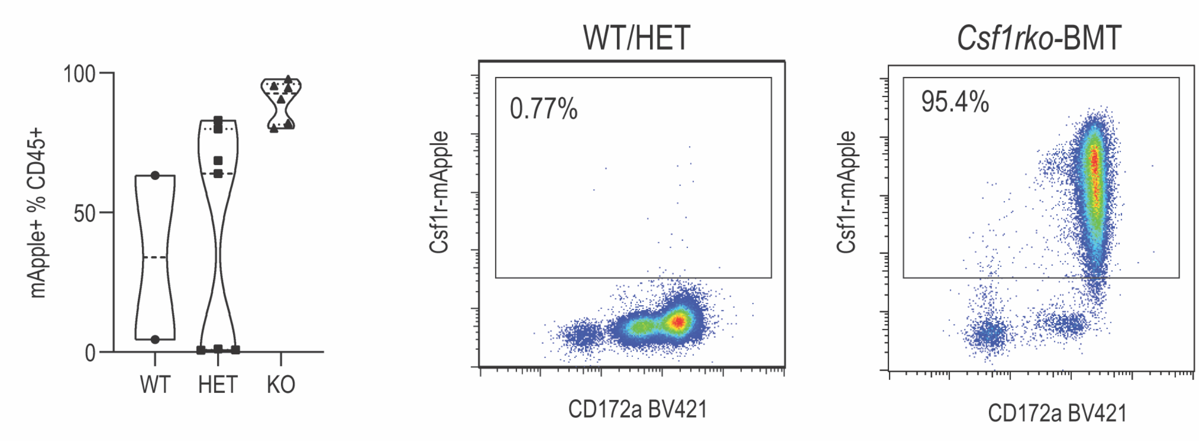


B. Peritoneal Csf1r-mApple+ cells 2-4 wks post-BMT

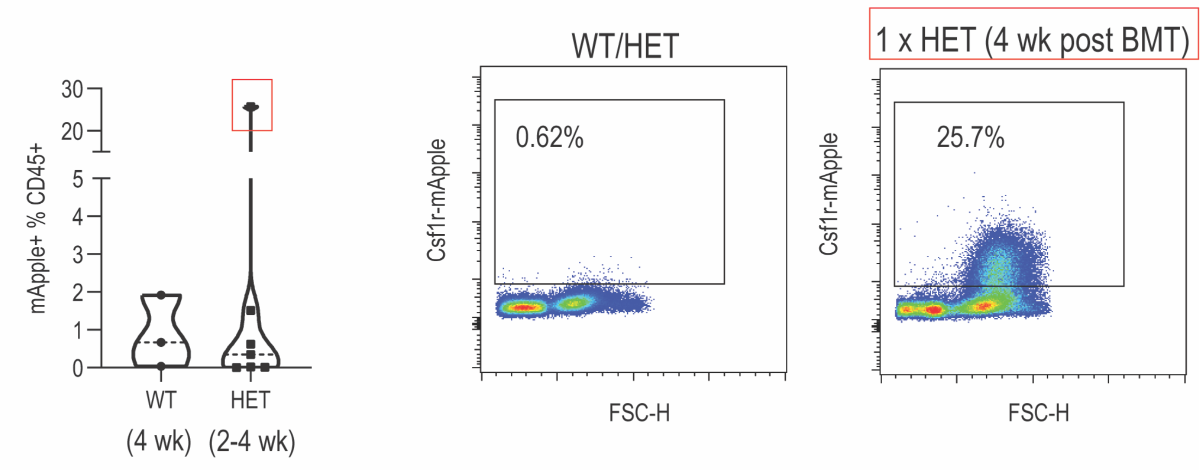

Figure S3

A. Csf1r-mApple+ cell profile in donor BM

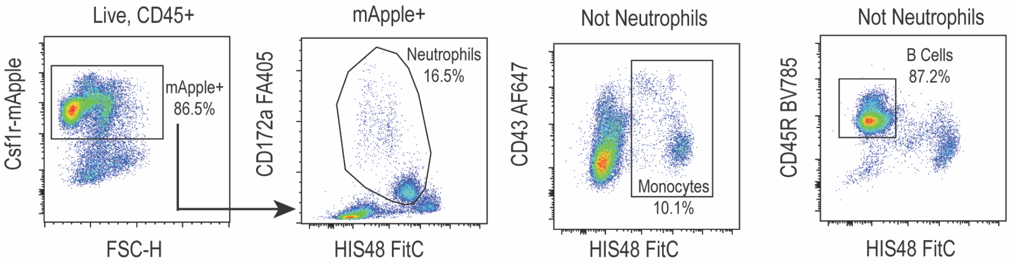


B. Csf1r-mApple+ cells in peripheral blood 1 wk post-BMT

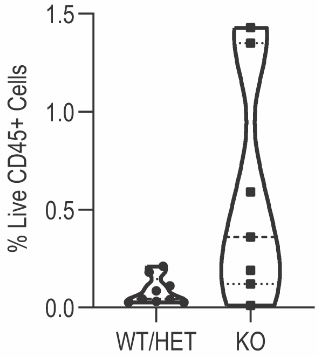


C. Csf1r-WT/HET Peripheral Blood 1 wk post BMT

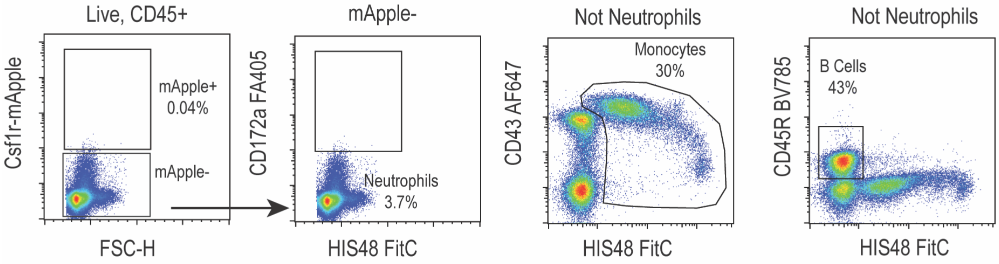


D. Csf1r-KO Peripheral Blood 1 wk post BMT

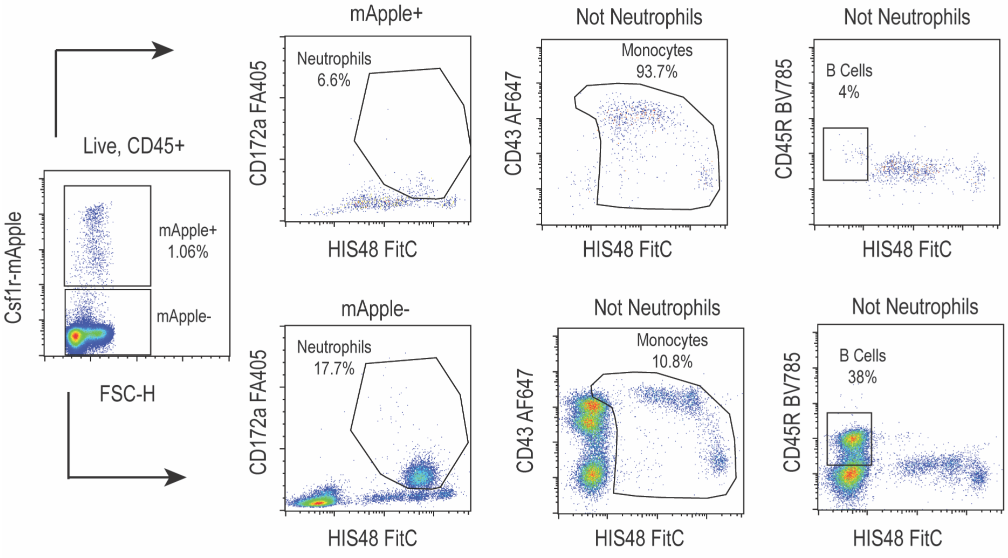
